# Horizontal acquisition of a patchwork Calvin cycle by symbiotic and free-living Campylobacterota (formerly Epsilonproteobacteria)

**DOI:** 10.1101/437616

**Authors:** Adrien Assié, Nikolaus Leisch, Dimitri V. Meier, Harald Gruber-Vodicka, Halina E. Tegetmeyer, Anke Meyerdirks, Manuel Kleiner, Tjorven Hinzke, Samantha Joye, Matthew Saxton, Nicole Dubilier, Jillian M. Petersen

**Affiliations:** Max Planck Institute for Marine Microbiology, Celsiusstrasse 1, D-28359 Bremen, Germany; Department of Geoscience, University of Calgary, Calgary, 2500 University Drive Northwest, Alberta T2N 1N4, Canada; Department of Plant and Microbial Biology, North Carolina State University, Raleigh 27695, North Carolina, USA; Department of Pharmaceutical Biotechnology, University of Greifswald, Institute of Pharmacy, Greifswald D-17489, Germany; Institute of Marine Biotechnology, Greifswald D-17489, Germany; Department of Marine Sciences, The University of Georgia, Room 159, Marine Sciences Bldg. Athens, GA 30602-3636; MARUM - Zentrum für Marine Umweltwissenschaften, University of Bremen, Leobener Str. 2, 28359 Bremen, Germany; Department of Microbiology and Ecosystem Science, Research Network Chemistry Meets Microbiology, University of Vienna, Althanstrasse 14, 1090 Vienna, Austria.; Center for Biotechnology, Bielefeld University, Universitaetsstrasse 27, 33615 Bielefeld, Germany

## Abstract

Although the majority of known autotrophs use the Calvin-Benson-Bassham (CBB) cycle for carbon fixation, all currently described autotrophs from the Campylobacterota (previously Epsilonproteobacteria) use the reductive tricarboxylic acid cycle (rTCA) instead. We discovered campylobacterotal epibionts (“*Candidatus* Thiobarba”) of deep-sea mussels that have acquired a complete CBB cycle and lost key genes of the rTCA cycle. Intriguingly, the phylogenies of campylobacterotal CBB genes suggest they were acquired in multiple transfers from Gammaproteobacteria closely related to sulfur-oxidizing endosymbionts associated with the mussels, as well as from Betaproteobacteria. We hypothesize that “*Ca.* Thiobarba” switched from the rTCA to a fully functional CBB cycle during its evolution, by acquiring genes from multiple sources, including co-occurring symbionts. We also found key CBB cycle genes in free-living Campylobacterota, suggesting that the CBB cycle may be more widespread in this phylum than previously known. Metatranscriptomics and metaproteomics confirmed high expression of CBB cycle genes in mussel-associated “*Ca.* Thiobarba”. Direct stable isotope fingerprinting showed that “*Ca.* Thiobarba” has typical CBB signatures, additional evidence that it uses this cycle for carbon fixation. Our discovery calls into question current assumptions about the distribution of carbon fixation pathways across the tree of life, and the interpretation of stable isotope measurements in the environment.

## Introduction

All life on earth is based on carbon fixation, and its molecular machinery is increasingly becoming a focus of biotechnology and geo-engineering efforts due to its potential to improve crop yields and sequester carbon dioxide from the atmosphere^1^. Seven carbon fixation pathways have evolved in nature, and one purely synthetic pathway runs in vitro^2–4^. Of the six natural pathways, the Calvin-Benson-Bassham (CBB) cycle was the first discovered, and is believed to be the most widespread^5–7^. The CBB cycle is used by a diverse array of organisms throughout the tree of life, including plants and algae, cyanobacteria, and autotrophic members of the Alpha-, Beta- and Gammaproteobacteria. Its key enzyme, the ribulose bisphosphate carboxylase/oxygenase (RuBisCO) is thought to be the most abundant, as well as one of the most ancient enzymes on Earth^8,9^.

The reductive tricarboxylic acid (rTCA) cycle was the second described carbon fixation pathway^10^. In short, it is a reversal of the energy-generating and oxidative TCA cycle. Instead of oxidizing acetyl-CoA and generating ATP and reducing equivalents, it reduces CO_2_ at the expense of ATP and reducing equivalents^2,7,10^. Most of the enzymes are shared with the TCA cycle, except for those that catalyze irreversible reactions in the TCA, such as citrate synthase, which is catalyzed by ATP citrate lyase in the rTCA. However, given sufficiently high reactant to product ratios and enzyme concentrations, the citrate synthase reaction can be reversed to run the TCA cycle reductively, without any additional enzymes^11,12^. The rTCA pathway is widely distributed in nature, and has been described in diverse lineages of anaerobes and microaerobes such as the Chlorobi, Aquificae, Nitrospirae and is also commonly observed among the Proteobacteria, including the Deltaproteobacteria and the Campylobacterota, (formerly Epsilonproteobacteria)^13,14^. It is particularly prominent in the Campylobacterota, as all previously described autotrophic members of this class use the rTCA pathway for CO_2_ fixation^2,13^.

Carbon fixation by chemoautotrophic microorganisms forms the basis of entire ecosystems at deep-sea hydrothermal vents and cold seeps^15,16^. Most of this carbon is fixed either via the CBB cycle, used by many gammaproteobacterial autotrophs, or the rTCA cycle, used by campylobacterotal autotrophs. This difference is reflected by the different niches colonized by these organisms at hydrothermal vents and seeps, with Gammaproteobacteria typically dominating habitats with higher oxygen and lower sulfide concentrations where the CBB cycle would be more efficient, and Campylobacterota typically thriving at lower oxygen and higher sulfide concentrations where the rTCA cycle could provide a selective advantage^17–23^. Experimental studies have linked substrate preferences in cultured Gammaproteobacteria and Campylobacterota to these ecological distributions^24–26^. Symbiotic invertebrates at hydrothermal vents and cold seeps associate with either gammaproteobacterial or campylobacterotal endosymbionts, which they rely on for most of their nutrition^27,28^. Some vent and seep invertebrates associate with both gammaproteobacterial and campylobacterotal symbionts simultaneously, which raises the question of how these co-occurring symbionts with differing habitat preferences can both be provided with suitable conditions^27,29,30^.

Bathymodiolin mussels, a subfamily of mytilid bivalves, are found at most hydrothermal vents and cold seeps^31^. They have evolved mutualistic relationships with chemosynthetic bacteria, allowing them to colonize these extreme environments. Inside their gills, they host intracellular sulfide- or methane-oxidizing gammaproteobacterial endosymbionts. Many bathymodiolin species host both types in a ‘dual symbiosis’. Assié et al. recently discovered a novel family of Campylobacterota, which colonizes bathymodiolin mussels from around the world^32^. In contrast to the gammaproteobacterial endosymbionts of these mussels that are harbored inside gill cells called bacteriocytes, these Campylobacterota are filamentous epibionts that colonize the surface of the gill epithelia in dense patches.

In this study, we used a multi-omics approach to investigate the metabolism of these novel epibionts in two different species of bathymodiolin mussels. Surprisingly, the epibionts have, and express, all the genes required for the CBB cycle but are missing key genes of the rTCA cycle. These CBB cycle genes were most likely acquired by horizontal gene transfer from diverse sources. With a recently developed, highly sensitive, direct stable isotope fingerprinting technique^33^, we show that the proteins of these epibionts have an isotopic signature typical of the CBB cycle, further demonstrating its importance for the metabolism of these epibionts. The discovery of Campylobacterota that employ the CBB cycle for CO_2_ fixation has wide-reaching implications for understanding the evolution of carbon fixation pathways, and for interpreting stable isotope values in environmental samples.

## Main text

### Genome assemblies and annotations

We assembled Campylobacterota draft genomes from gill metagenomes of two mussel species: *“B.” childressi* and *B. azoricus*. The draft genome from *“B.” childressi* was 2.2 Mb, and estimated to be 95% complete. It had 30% GC, 2204 predicted protein-coding genes and 31 tRNA-encoding genes. The draft genome from *B. azoricus* was estimated to be 92% complete at 2.3 Mb. It had 30% GC, 2155 predicted protein-coding genes and 37 tRNAs (Table S1). The draft genomes had an average nucleotide sequence identity (ANI) of 83.1%, indicating that they represent different species belonging to the same genus^34–37^.

Previous 16S ribosomal RNA gene (16S rRNA) sequence analysis identified this group of epibionts as a novel family-level deep-branching sister group of the *Sulfurovum* clade within the Campylobacterota^32^. The Campylobacterota draft genome from *“B.” childressi* contained a partial 16S rRNA sequence (586 bp) that was 100% identical to the epibiont sequence previously published^32^. The 16S rRNA sequence identity and the ANI information allowed us to link the two draft genomes to the previously described epibiont^32^. To better resolve the relationships of the mussel epibionts to other Campylobacterota we analyzed a set of 18 conserved marker genes from the two epibiont draft genomes and other publicly available Campylobacterota genomes (Figure S1). In contrast to the previous 16S rRNA based phylogeny^32^, our analysis placed the mussel epibionts on a long branch, basal to the main Campylobacterota families. The long-branch formation for the genomes presented in this study is likely related to low amino acid sequence identity (AAI) values between these and the Campylobacterota representative genomes. AAI values were below 48% when comparing the Campylobacterota bins found in our bathymodiolin samples with their closest relative genomes, *Sulfurospirillum arcachonense* and *Arcobacter anaerophilus* (Supplementary Table 2). According to the guidelines of Rodriguez and Konstantinidis^36^, organisms with AAI values higher than 30% and lower than 55-60% are likely to belong to the same division, but not the same genus. This is corroborated on the 16S rRNA level^37^, where our genomic data supports the previously published 16S rRNA based study^32^, indicating that the epibiont species belong to a novel family of Campylobacterota.

We therefore propose the new *Candidatus* family “Thiobarbaceae” (Campylobacterales, Campylobacterota), with the name composed of “*Thio-*” from the Greek word θ∊ῖον, theîon for sulfur and “*barba*” from the Latin word for beard. The proposed family includes the novel *Candidatus* genus “Thiobarba” with two *Candidatus* species “*Ca.* T. azoricus” and “*Ca.* T. childressi”, for the two epibiont species in reference to their respective hosts, *B. azoricus* and *B. childressi*. For more details on the aetiology see Supplementary note 1.

### Unexpected carbon fixation pathways of “*Candidatus* Thiobarba spp.”

Considering their phylogenetic relationship to free-living chemolithoautotrophic and mixotrophic Campylobacterota and their presence in sulfide-rich environments, we searched the epibiont draft genomes for metabolic pathways indicative of heterotrophy, autotrophy and sulfur oxidation. Both “*Ca.* Thiobarba” genomes encoded all the genes for the SOX multi-enzyme pathway of sulfur oxidation, and are thus capable of lithotrophy using reduced sulfur compounds as electron donors (Figure 1). Like other sulfur-oxidizing Campylobacterota, they also appear capable of heterotrophic growth as their genomes contained a TCA cycle and a partial glycogenesis/glycolysis pathway (Supplementary note 2 and Figure S2).

**Figure 1.**
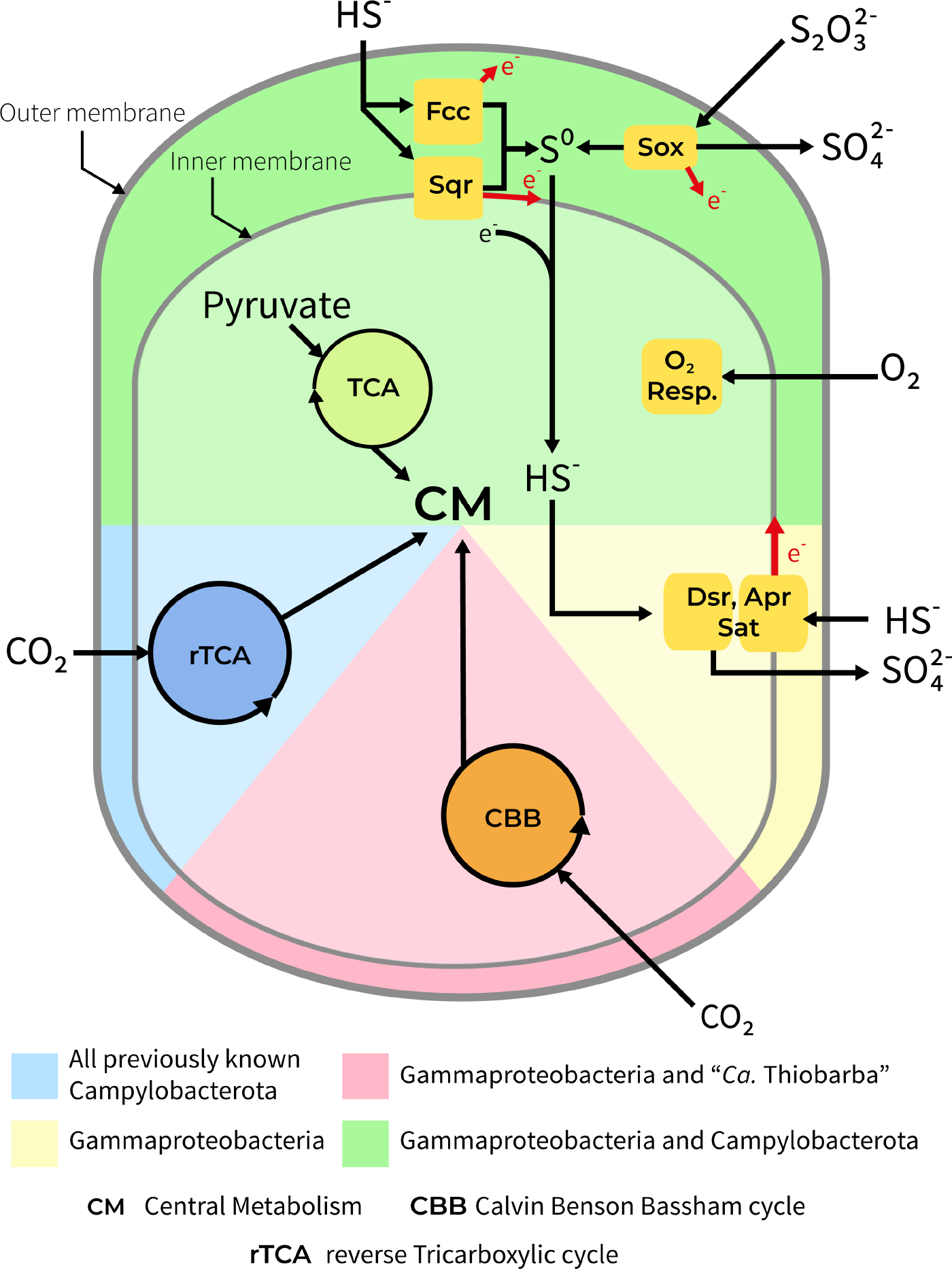
“*Ca*. Thiobarba spp.” share metabolic features of Gammaproteobacteria and Campylobacterota. Figure shows overview of the main metabolic pathways for energy generation and carbon fixation in known chemosynthetic Gammaproteobacteria and Campylobacterota compared to the metabolism of “*Ca.* Thiobarba spp.”.

All previously described sulfur-oxidizing Campylobacterota use the reverse TCA cycle for carbon fixation^2^. All of these bacteria have genes encoding the enzymes for this cycle including the pyruvate: ferredoxin oxidoreductase genes *porABCD*, the 2-oxoglutarate oxidoreductase genes *oorABDG*, and the ATP citrate lyase genes *aclAB*. Unexpectedly, we could not find most of these genes in the “*Ca.* Thiobarba” genomes. The “*Ca.* T. childressi” draft genome contained only the *porAB* genes, and the “*Ca* T. azoricus” draft genome contained *porABCD* and *aclA*, but not the *oorABDG* genes. To confirm that these genes were not missing because of errors in assembly, binning or annotation, we searched the genomes and unbinned metagenome assemblies with BLAST. No additional rTCA cycle genes could be found in the draft genomes or in the entire *“B.” childressi* metagenome assembly (Supplementary note 3). The absence of rTCA cycle genes suggests that either a) the epibionts never had a complete rTCA cycle, or b) it has been lost over the course of evolution. The additional roles of the *por* genes in other metabolic pathways, such as pyruvate fermentation, could explain why these are present, at least in part, in both lineages^38^.

Although the rTCA cycles were incomplete, both “*Ca.* Thiobarba” genomes contained all the genes required for carbon fixation via the CBB cycle (Figure 1). Most CBB cycle enzymes are used in other metabolic pathways, and are thus also found in heterotrophic bacteria, but two enzymes are unique to the cycle: Phosphoribulokinase (PRK) and ribulose 1,5-bisphosphate carboxylase/oxygenase (RuBisCO)^2^. In both “*Ca.* Thiobarba” species, 9 out of the 12 genes encoding PRK, RuBisCO and accessory proteins were grouped in two clusters, while three additional genes for the CBB cycle were scattered on separate contigs (Figure S3). The first cluster consisted of the RuBisCO Form I large and small subunits (*rbcL* and *rbcS*), a conserved hypothetical protein, and the RuBisCO activation protein *cbbQ.* The order of these genes was conserved in both epibionts (Figure S4). “*Ca.* T. childressi” had an additional gene encoding the RuBisCO activation protein *cbbO* in this first cluster. In the “*Ca.* T. azoricus” genome, this gene was located on a separate contig. The second cluster included the genes coding for fructose-1,6-bisphosphatase, PRK, transketolase, phosphoglycolate phosphatase (not known to be involved in the CBB cycle), fructose-bisphosphate aldolase and ribulose-phosphate 3-epimerase (Figure S3). The order of the second gene cluster was consistent in both epibiont genomes, but the gene neighborhoods surrounding this cluster differed (Figure S4).

### CBB cycle expression in “*Ca*. Thiobarba childressi”

To confirm expression of the CBB cycle by the epibionts, we analyzed the metatranscriptomes and -proteomes of *“B.” childressi*, the mussel species with the highest abundance of these epibionts^32^. We found that all CBB cycle genes were expressed in the transcriptomes, including the *rbcL* and *rbcS*, which were among the most highly expressed genes of this epibiont (Table 1). Although “*Ca* T. childressi” was present in relatively low abundance in the metaproteome samples (~0.5% of the total sample proteinaceous biomass, calculated according to ^39^), the RuBisCO small and large subunits were among the unique “*Ca.* T. childressi” proteins detected, further indicating high expression levels. The abundance of CBB cycle transcripts and proteins highlights their importance in the metabolism of “*Ca.* T. childressi” (for full transcription and expression information, see Tables S3 and S4).

**Table 1.** Transcription and translation ranks for the detectable genes involved in the CBB cycle of “*Ca.* T. childressi”.

| Gene_ID | Name | Transcription<br>rank | Translation<br>rank |
| --- | --- | --- | --- |
| **BCM6EPS_1532** | Ribulose biphosphate carboxylase large chain<br>EC 4.1.1.39 | 14 | 18 |
| **BCM6EPS_1531** | Ribulose biphosphate carboxylase small chain<br>EC 4.1.1.39 | 17 | 19 |
| **BCM6EPS_1455** | NAD-dependent glyceraldehyde-3-phosphate<br>dehydrogenase<br>EC 1.2.1.12 | 75 | Not detected |
| **BCM6EPS_1028** | Transketolase<br>EC 2.2.1.1 | 80 | Not detected |
| **BCM6EPS_1030** | Fructose-bisphosphate aldolase class II<br>EC 4.1.2.13 | 90 | Not detected |
| **BCM6EPS_1031** | Ribulose-phosphate 3-epimerase<br>EC 5.1.3.1 | 150 | Not detected |
| **BCM6EPS_1027** | Phosphoribulokinase<br>EC 2.7.1.19 | 160 | Not detected |
| **BCM6EPS_1026** | Fructose-1-6-bisphosphatase- type I<br>EC 3.1.3.11 | 182 | Not detected |
| **BCM6EPS_1029** | Phosphoglycolate phosphatase<br>EC 3.1.3.18 | 236 | Not detected |
| **BCM6EPS_1456** | Phosphoglycerate kinase<br>EC 2.7.2.3 | 440 | Not detected |
| **BCM6EPS_514** | Triosephosphate isomerase<br>EC 5.3.1.1 | 461 | Not detected |

### Direct stable isotope fingerprinting confirmed a CBB signature for “*Ca*. Thiobarba childressi”

The stable carbon isotope signatures of an environmental sample reflect the pathway that dominates inorganic carbon fixation in the chemoautotrophic members of the bacterial community^40^. Due to differences in kinetic isotope effects, the enzymes involved in the different carbon fixation pathways vary in the degree to which they discriminate against the heavier ^13^C. This leads to a shift in the ^12^C/^13^C ratio between the inorganic carbon source and the generated biomass that is characteristic for the carbon fixation pathway. The CBB cycle generates a −13 to −26‰ shift of the δ^13^C ratio, while the rTCA cycle leads to a much smaller −3 to −13‰ shift^40^.

The average δ^13^C value of bulk *“B.” childressi* gill tissues was −47.1 ± 2.6‰, (Table S5). However, these values reflect the stable isotope composition of all members of the symbiotic community. As most of the biomass is from the host animal or the highly abundant methane-oxidizing gammaproteobacterial endosymbiont, the signal of the epibiont is greatly diluted^41^. To overcome this limitation and to distinguish between the stable carbon isotope values of the symbiotic partners, we employed the recently-developed direct Protein-SIF method (SIF = stable isotope fingerprinting) on our metaproteomic data set^33^. Direct Protein-SIF quantifies the stable isotopic composition of uncultivated members of a mixed community for which genomes or transcriptomes are available. Peptides from the methane-oxidizing symbionts had a δ^13^C of −38.8 ± 0.7‰, and host peptides had −44.2 ± 0.6‰. These values are similar to those of the methane gas at this cold seep site. Thus, the methane-oxidizing symbionts likely obtain most of their carbon from methane^42,43^. The host values were similar to those of bulk measurements. However, they were unexpectedly light compared to the methane-oxidizing symbionts, considering that these mussels are thought to gain most of their nutrition from their methane-oxidizing symbionts, and would therefore be expected to have similar δ^13^C values. As *B. childressi* is known to be capable of filter-feeding^44^, these values possibly reflect nutritional supplementation from filter-feeding on microorganisms with even lighter δ^13^C values than the methane-oxidizing symbionts, that is from the seep environment (as phototrophic microorganisms from the surface would have heavier δ^13^C values).

“*Ca.* T. childressi” had a much lower abundance in the metaproteomic dataset compared to the host and the methane-oxidizing symbionts. Nevertheless, we were able to detect 50 peptides that were unique to “*Ca.* T. childressi”. This allowed us to estimate its natural δ^13^C value, which was relatively light at −66.6 ± 12.5‰. There are two possible inorganic carbon sources for these epibionts: 1) ambient seawater inorganic carbon, which has a δ^13^C value of +3‰^42^, and 2) inorganic carbon produced as an end product of methane oxidation by methane-oxidizing bacteria or respiration by the host, which we expect to be around −39‰ for a gas hydrate site, similar to our collection site^41,42^. We calculated the expected values of biomass generated if either of these carbon sources were fixed through the rTCA cycle or the CBB cycle (Figure 2). Regardless of the inorganic carbon source, the δ^13^C values of “*Ca.* T. childressi” peptides are far lighter than would be expected if they used the rTCA cycle. They are, however, consistent with the values for carbon fixation that could be expected when “*Ca.* T. childressi” used the CBB cycle, with inorganic carbon derived from symbiont methane oxidation or host respiration.

**Figure 2.**
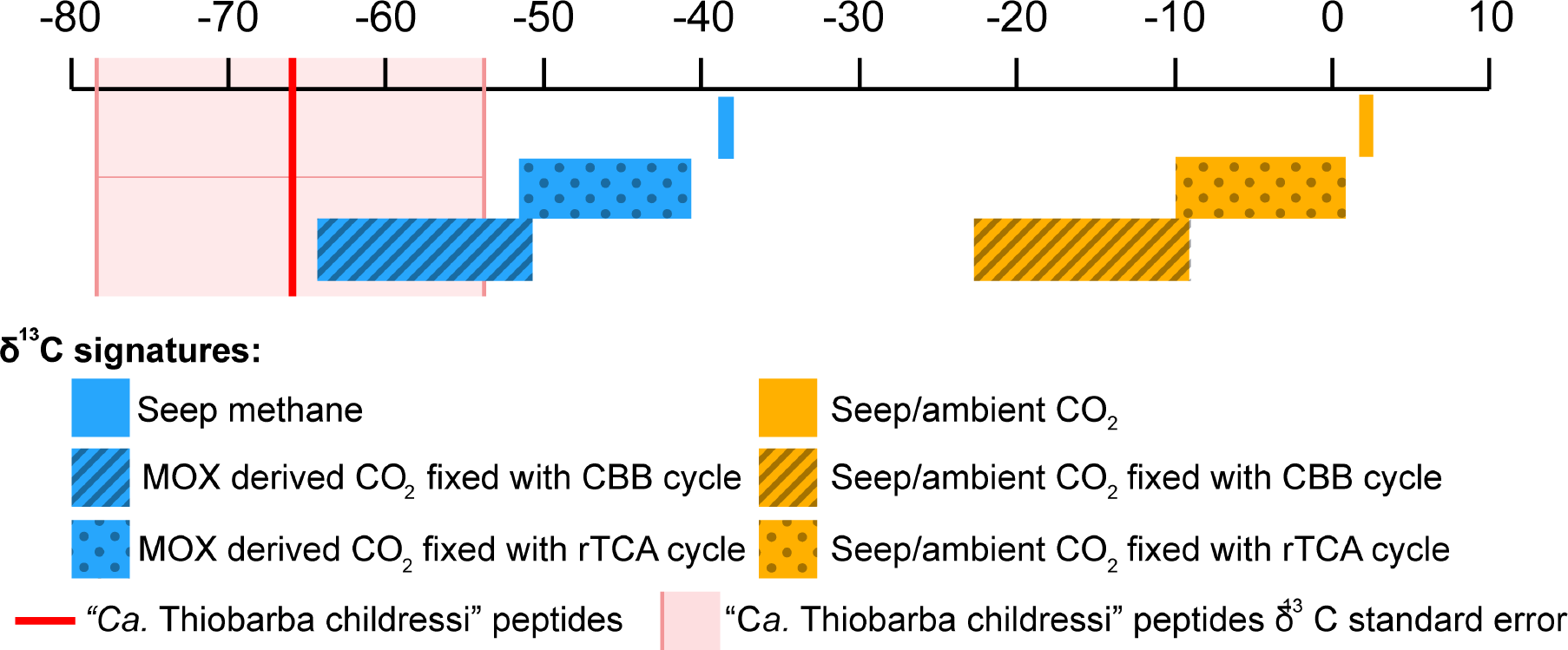
Stable carbon isotope values of “*Ca*. Thiobarba childressi” are consistent with carbon fixation via the CBB cycle. Model of δ^13^C values of deep-sea carbon and the predicted influence of different inorganic fixation pathways on these values. The δ^13^C values of CO_2_ originating from ambient seawater are shown in yellow, and the expected δ^13^C values of CO_2_ originating from methane oxidation are shown in blue. The red line represents the average δ^13^C value measured for “*Ca.* Thiobarba” peptides using direct Protein-SIF. Reference δ^13^C values for “Seep methane” and “Seep/ambient CO_2_“ are based on Macavoy et al.^41^ and Sassen et al.^42^. Transformations of δ^13^C values for each metabolic pathway are estimated based on Pearson et al.^40^.

### Calvin cycle genes in free-living Campylobacterota

After discovering the CBB cycle in the mussel epibionts, we asked if other members of the Campylobacterota might also have acquired these genes. We discovered key CBB cycle genes in a Campylobacterota draft genome binned from a metagenomic library from diffuse hydrothermal fluids collected in the Manus Basin (Western Pacific)^23^. This draft genome was composed of 60 contigs with 29.1% GC content, and based on the CheckM single-copy genes set, was 92.2% complete^45^. Phylogenomic reconstruction placed this organism on a deep branch basal to the Arcobacteraceae family. AAI values showed between 55 and 58% similarity with the Arcobacteraceae, thus, this Campylobacterota bin might belong to a new genus within the Arcobacteraceae (Supplementary Table 2). Our phylogenetic analyses and calculated AAI values clearly show that this environmental bin belongs to a Campylobacteraota family distinct from “*Ca.* Thiobarba” (Figure S1). Although a full rTCA cycle was present in the draft genome, we also found genes coding for a RuBisCO form I enzyme, a hypothetical gene and CbbQ in one cluster. This cluster shared the same gene order, as well as 84% nucleotide sequence identity, with the CBB cycle cluster we found in “*Ca.* Thiobarba” (Figure S4). The high sequence similarity between these clusters suggests a similar origin for both of them. If these genes and the enzymes they encode are active in the Manus Basin organism, then free-living Campylobacterota may also be able to use the CBB cycle to fix carbon. These bacteria could be using both cycles depending on the environmental setting, as suggested for the sulfide-oxidizing gammaproteobacterial symbionts of vestimentiferan tubeworms^46–48^ found at hydrothermal vents, the large sulfur bacteria *Beggiatoa* and *Thiomargarita* spp.^49–51^, and recently the cultivable sulfur oxidizer *Thioflavicoccus mobilis*^52^. The tubeworm symbiont *Ca.* E. persephone expresses both the CBB and the rTCA cycle in the same host individual, but it is still unclear how these two cycles are coordinated at the level of individual symbiont cells, or over time^47,48^.

### Possible evolutionary origins of Campylobacterota CBB genes

Considering the lack of CBB cycle genes in all Campylobacterota investigated prior to this study, it is most likely that this carbon fixation pathway was acquired by “*Ca.* Thiobarba” and free-living Campylobacterota through horizontal gene transfer, rather than being an ancestral pathway in this phylum. We investigated the evolutionary origins of the genes coding for CBB enzymes, including those with additional roles in other metabolic pathways, using BLAST analyses of nucleotide and protein sequences, and phylogenetic reconstruction of protein sequences. BLAST analyses revealed that only two of the “*Ca.* Thiobarba” CBB genes were affiliated with genes from other Campylobacterota. Of the other 10, five had best hits to Gammaproteobacteria, and five had best hits to Betaproteobacteria (Table S6).

Phylogenetic reconstruction further supported our hypothesis that the “*Ca.* Thiobarba” CBB cycle is a ‘patchwork’ of genes with evolutionary origins in the Betaproteobacteria, Gammaproteobacteria, and Campylobacterota (Figure 3). The RuBisCO large and small subunits *rbcL* and *rbcS*, their accessory proteins *cbbQ* and *cbbO*, as well as the glyceraldehyde-3-phosphate dehydrogenase proteins clustered with a clade of gammaproteobacterial sulfur-oxidizing chemolithoautotrophs. Many of the related sequences belonged to free-living sulfur oxidizers such as “*Ca.* Thioglobus autotrophicus” and the gammaproteobacterial sulfur-oxidizing endosymbionts of bathymodiolin mussels (Figure 4). Phylogenetic analysis of “*Ca.* Thiobarba” PRK proteins placed these on a long branch between gamma-, alpha- and betaproteobacterial clades, but this placement did not have high support (Figure 5). This could indicate that the “*Ca*. Thiobarba” PRK proteins truly belong to a Campylobacterota gene family, and because these are the first sequences available from this family, their phylogenetic placement is currently not well supported.

**Figure 3.**
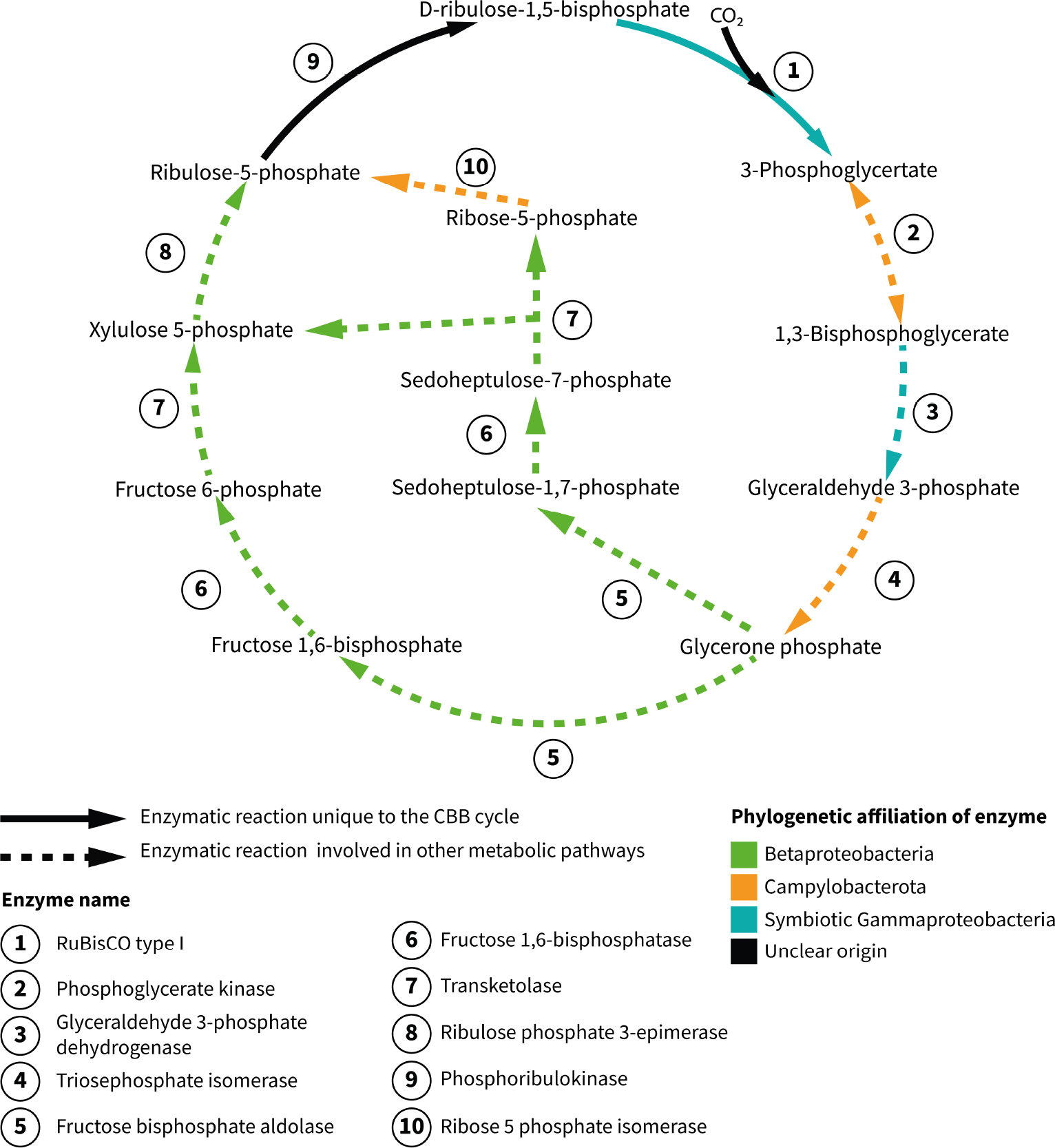
“*Ca*. Thiobarba” genomes encode a CBB cycle with genes affiliated to at least three phylogenetically distinct classes. The solid arrows indicate enzymatic reactions that are unique to the CBB cycle, while the dashed arrows indicate that the enzymes are also involved in other metabolic pathways. Enzyme names are shown in bold and the colors represent their phylogenetic affiliations.

**Figure 4.**
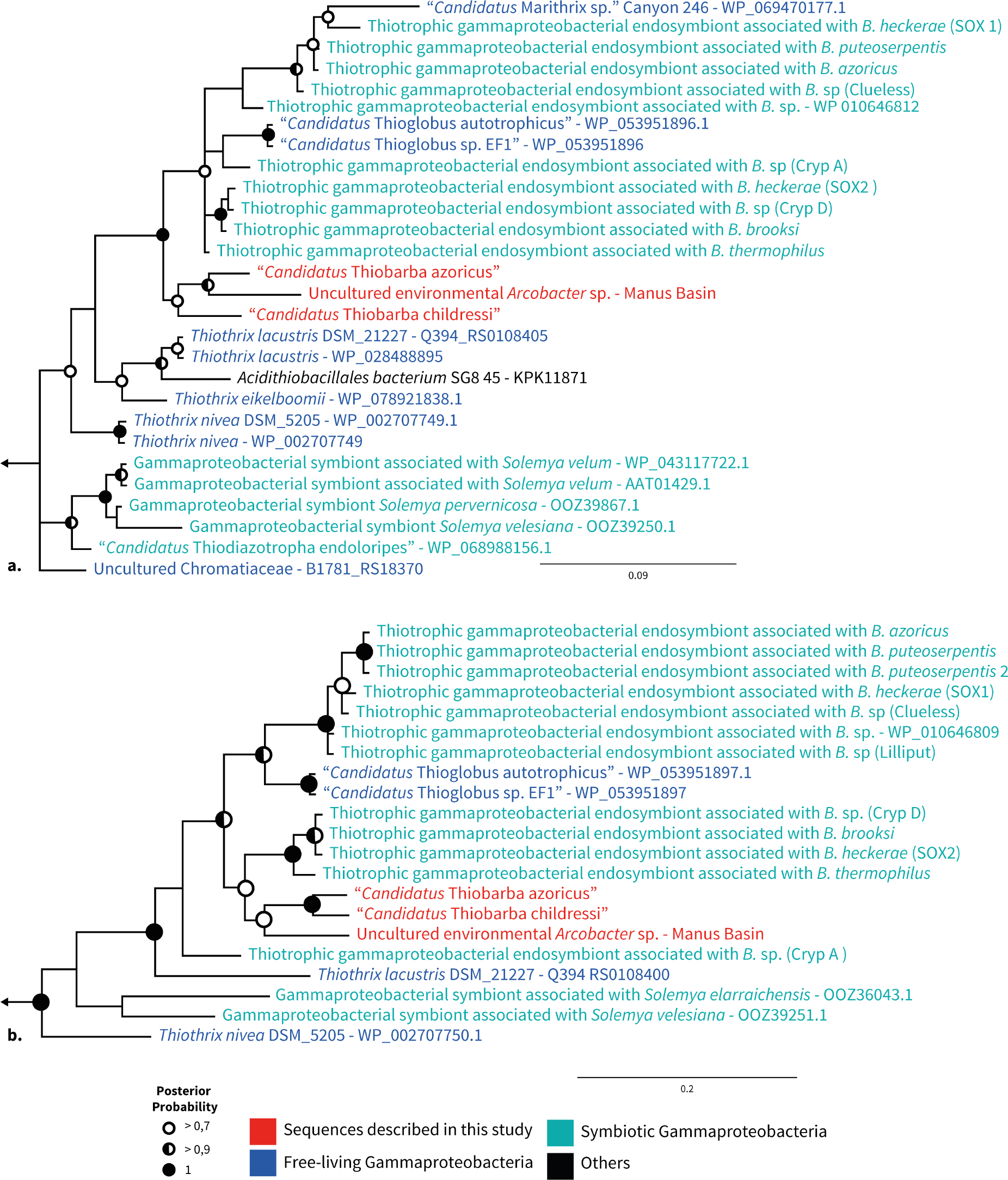
“*Ca*. Thiobarba” RuBisCO proteins cluster with gammaproteobacterial sequences. Bayesian inference trees of RuBisCO large (a) and small (b) subunit amino acid sequences under an LG model with Gamma-distributed rates of evolution. Analyses were performed with 6 million generations using two parallel Monte Carlo Markov chains. Sample trees were taken every 25000 generations. Left arrows indicate truncated tree, tree roots were built from *Prochlorococcus* and *Synechococcus* sequences for (a) and *Planktothrix* and *Synechococcus* sequences for (b). Full trees are displayed as Supplementary figures 14 and 15.

**Figure 5.**
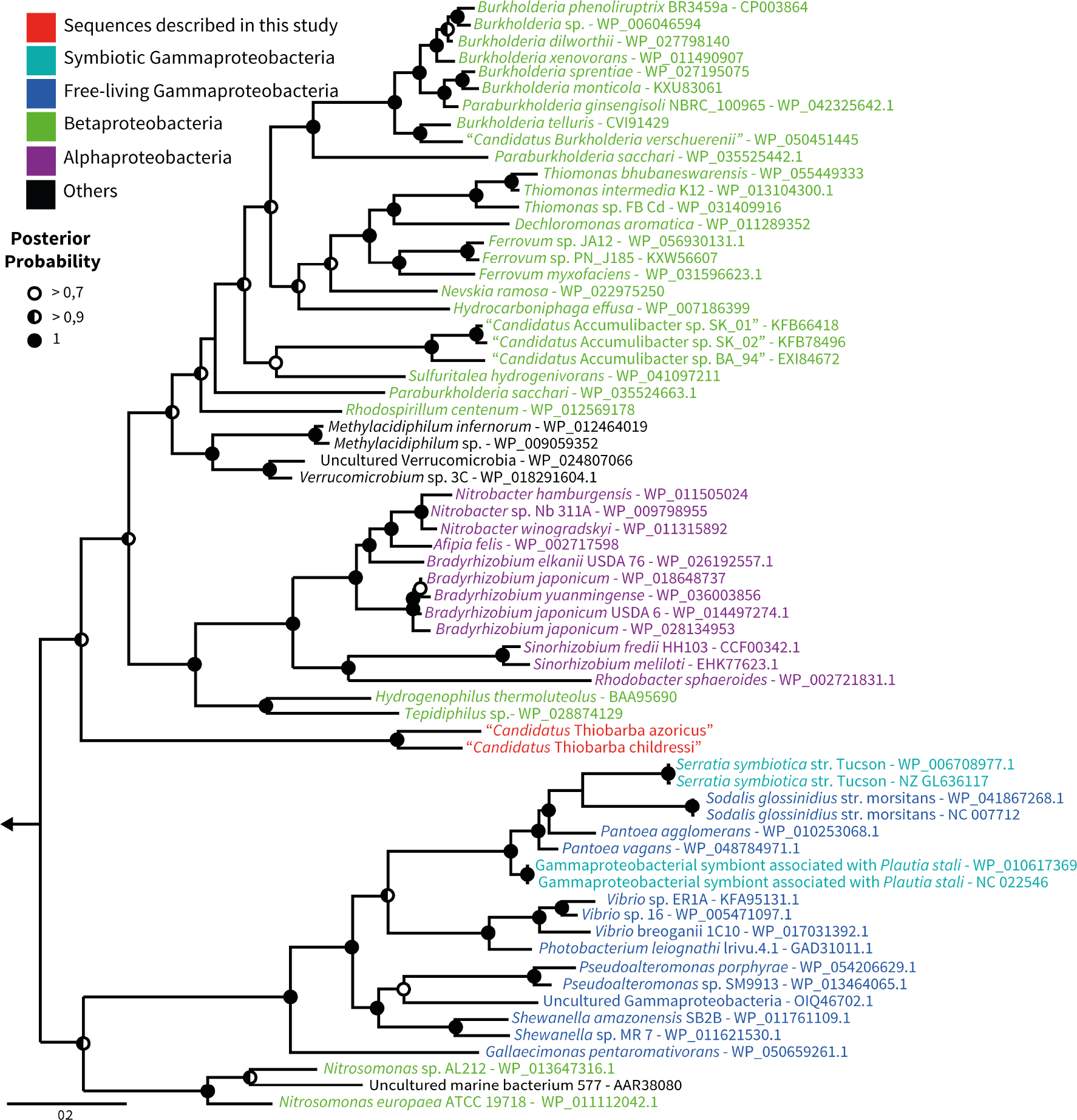
“*Ca*. Thiobarba” phosphoribulokinases are loosely affiliated with those from Betaproteobacteria, Alphaproteobacteria, and Verrucomicrobia. Bayesian inference tree of phosphoribulokinase amino acid sequences under an LG model with Gamma-distributed rates of evolution and a proportion of invariant sites. Analyses were performed with 6 million generations using two parallel Monte Carlo Markov chains. Sample trees were taken every 25000 generations. Left arrow indicates truncated root, the root is built from distant *Prochlorococcus* and *Synechococcus* sequences. Full tree is displayed as supplementary figure 23.

Further sampling may help to clarify their evolutionary history. Four “*Ca.* Thiobarba” CBB cycle proteins consistently belonged to a sister branch to betaproteobacterial sequences (fructose 1,6-bisphophatase, 1,6-bisphophate aldolase, transketolase and ribulose phosphate 3-epimerase). Only two proteins were phylogenetically related to those from other Campylobacterota (phosphoglycerate kinase and triose phosphate isomerase) (Figures S5 to S24).

The CBB genes consistently fell into three phylogenetic groups: some CBB genes were most closely related to those of other Campylobacterota, some to those of Gammaproteobacteria, and some to those of Betaproteobacteria. The “*Ca*. Thiobarba” CBB genes that fell within the Gammaproteobacteria were similarly organized to genes with which they were most closely related, such as those from “*Ca.* Thioglobus autotrophicus” and the endosymbionts of bathymodiolin mussels (Figure S24 and Supplementary note 4). Similarly, the genes that were most closely related to Betaproteobacteria had a similar organization to genes found in free-living Betaproteobacteria such as *Paraburkholderia xenovorans* (NC_007651) and *Dechloromonas aromatica* (NC_007298) (Figure S24). This similarity further supports the hypothesis that these CBB cycle genes were acquired by “*Ca*. Thiobarba” at least twice, in independent horizontal gene transfer (HGT) events, with one possibly originating from Gammaproteobacteria, and another possibly from Betaproteobacteria. Alternatively, it is also possible that the Betaproteobacteria-like genes clustering on long branches, such as the PRK, are Campylobacterota genes that have not previously been sequenced. Regardless of the number of HGT events, the acquisition of these genes presumably happened in a common ancestor to the “*Ca.* Thiobarbaceae”. Codon usage analysis supports our hypothesis of a relatively ancient acquisition, as the codon usage of the CBB genes was similar to that of the “*Ca.* Thiobarbaceae” core genome (Figure S25). After horizontal acquisition, the codon usage initially reflects the original donor’s, but over time mutates to match its host’s^53^.

Gene order within each of the two CBB clusters was identical in “*Ca.* T. azoricus” and “*Ca.* T. childressi”. This further supports our hypothesis of a single acquisition event for each cluster in a common ancient ancestor. This synteny also highlights the tendency of these clusters to resist genomic rearrangements. In contrast, the genomic neighborhoods of the CBB clusters differed between the two “*Ca.* Thiobarba”, indicating that subsequent genome rearrangements occurred since the divergence of these two epibionts. Mobile element genes and transposases were the most highly expressed genes in “*Ca.* T. childressi” based on our transcriptomes, which, if active, could explain these rearrangements (Table S3)^54^.

### Evolutionary advantages of the CBB cycle

Members of the Campylobacterota occupy remarkably diverse habitats, and have a range of different lifestyles and metabolic capabilities, from chemolithoautotrophs that use a suite of electron donors and acceptors, to heterotrophic symbionts and pathogens of humans and other animals^13,55^. Evolutionary studies suggest that Campylobacterota emerged in deep-sea habitats, subsequently colonizing and diversifying across terrestrial and human-associated environments^13,56^. Considering the distribution of chemosynthetic potential within the Campylobacterota, it has been hypothesized that they evolved from an autotrophic common ancestor that first used the Wood-Ljungdahl pathway before switching to a more flexible rTCA cycle^13,17,57^. We hypothesize that in the symbiotic “*Ca*. Thiobarba” lineage, the rTCA cycle was replaced by yet another carbon fixation pathway, the CBB cycle.

Several environmental and genomic factors provide important clues as to why the CBB cycle was selected over the rTCA cycle in “*Ca*. Thiobarba”. Both the CBB and rTCA cycles serve the same purpose, the fixation of inorganic carbon to provide building blocks for cell biomass. But a major difference between the two known carbon fixation pathways is their energy requirements. For example, if one molecule of pyruvate is synthesized from CO_2_ via the CBB cycle seven molecules of ATP are used, while the rTCA cycle only requires two ATP^2^. From an evolutionary point of view, exchanging a more energy-efficient carbon fixation pathway with a costlier one could best be explained if it comes with an additional advantage such as oxygen tolerance. The rTCA cycle relies on ferredoxin-based enzymes, which are quickly oxidized by oxygen, and as a result, most organisms with an rTCA cycle are anaerobes or microaerobes^58,59^. In contrast, CBB cycle enzymes are less affected by oxygen^60^. “*Ca.* Thiobarba” species colonize the gills of bathymodiolin mussels, a gas exchange organ that is exposed to oxygen and is typically dominated by gammaproteobacterial endosymbionts. The close phylogenetic relationship between some “*Ca.* Thiobarba” CBB genes with those from the sulfur-oxidizing gammaproteobacterial endosymbionts of bathymodiolin mussels suggests that either i) both symbionts acquired CBB genes from the same source or ii) “*Ca.* Thiobarba” acquired key genes from the gammaproteobacterial endosymbionts already adapted to the mussel gill niche.

Many deep-sea Campylobacterota grow attached to surfaces^29,61,62^, thus, a “*Ca.* Thiobarba” ancestor might have colonized mussel gills prior to acquiring the CBB cycle. Living attached to the gills would bring these epibionts into close proximity to the gammaproteobacterial endosymbionts. Sharing a niche has been shown to be a stronger predictor of horizontal gene transfer than phylogenetic relatedness^63^. Moreover, Campylobacterota have remarkably flexible genomes, with rampant genomic rearrangement and DNA uptake^64–66^. This affinity for foreign DNA uptake, and the physical proximity of epibionts and endosymbionts support scenario ii) above. The acquisition of the CBB carbon fixation pathway may have enabled “*Ca.* Thiobarba” to thrive attached to an animal host, leading to the complete reliance on the CBB cycle for carbon fixation and the gradual loss of the rTCA cycle.

Another major difference between the CBB and rTCA cycles is the metabolic end product: Sugar precursors such as glyceraldehyde 3-phosphate for the CBB cycle, and acetyl-CoA for the rTCA cycle^2,60^. Intriguingly, the “*Ca.* Thiobarba” genomes encoded many pathways requiring sugars, including N-linked glycosylation, capsular polysaccharides and lipooligosaccharide synthesis, some of which are not found in related chemolithoautotrophic Campylobacterota, but are present in heterotrophic host-associated Campylobacterota. “Ca. Thiobarba’s” ability to gain sugar precursors while fixing CO_2_ using the CBB cycle might be advantageous, despite the higher energy requirements compared to the rTCA cycle. Many of the pathways requiring sugars are predicted to play a role in surface structures and extracellular polysaccharide capsule formation, which can be key mediators of host attachment, and thus may be essential for its epibiotic lifestyle^67,68^.

### Evolving a Calvin cycle in nature and the laboratory

The complex metabolic network that links carbon fixation and central carbon metabolism poses a massive challenge to switching carbon fixation pathways, either in nature or in the laboratory. These links are usually specific to each pathway and to each organism^60^. Efforts to introduce non-native carbon fixation pathways have mainly focused on the CBB cycle because theoretically, only two additional enzymes are needed to run this cycle, even in heterotrophs such as *E. coli*^69,70^. However, a number of challenges must be overcome to express ‘foreign’ carbon fixation pathways in new organisms. In addition to the challenges inherent in expressing horizontally acquired genes, such as non-native promoter and codon usage, and the need for chaperones and biosynthesis enzymes, gene expression must be tightly regulated to balance the production and consumption of intermediates and end products. Because of this, to run the CBB cycle in engineered *E. coli*, the CBB cycle had to be synthetically decoupled from gluconeogenesis by deleting the phosphoglycerate mutase gene^69^. Switching from one carbon fixation pathway to another may be simpler in chemolithoautotrophs than re-wiring a chemoorganoheterotroph such as *E. coli* to use the CBB cycle. In a chemolithoautotroph, production of energy and reducing equivalents are already decoupled from carbon fixation, as they are generated through oxidation of reduced compounds such as sulfur. Nevertheless, switching from the rTCA to the CBB cycle is a major shift in cellular metabolism, requiring adaptation of diverse biosynthetic pathways linked to carbon fixation. As far as we are aware, this has not yet been observed in nature, but in the laboratory, *E. coli* required extensive fine-tuning of metabolic enzymes beyond the CBB cycle through experimental evolution to run a fully functional CBB cycle^3,69^.

## Conclusions

The environment is a potent driving force in structuring symbiotic and free-living microbial communities^27,30,71^. The distribution of gammaproteobacterial and campylobacterotal sulfur oxidizers is a typical example of adaptation to a geochemical niche; in a range of environments from hydrothermal vents^23^ and cold seeps^21^ to oxygen minimum zones^72^ and coastal sediments^73^, Gammaproteobacteria are usually associated with low-sulfide, high-oxygen environments, and campylobacterota with high-sulfide, low-oxygen environments. The horizontal acquisition of the CBB cycle genes may have allowed campylobacterotal “*Ca.* Thiobarba” to establish a symbiotic relationship in a niche that is usually dominated by Gammaproteobacteria.

The diverse origins of “*Ca*. Thiobarba’s” CBB cycle genes showcases the modularity^74^ of bacterial metabolism and demonstrates that in principle, fully functional metabolic cycles can be pieced together with enzymes from different organisms, both in the laboratory^3^ and in nature. In addition to acquiring the two genes theoretically required by a heterotroph to encode a full CBB cycle, “*Ca*. Thiobarba” seems to have replaced an extensive set of additional CBB genes. This suggests that similar to laboratory models, this natural metabolic switch required ‘tweaking’ of further enzymes of this pathway, and possibly other pathways that siphon off intermediates. Metabolic modularity is considered one of the main factors organizing biological networks^74^. Understanding genome evolution in “*Ca.* Thiobarba” will shed light on the complex interplay between gene acquisition, expression and the selection that caused the evolution of this major metabolic shift. Our findings highlight the central role that horizontal gene transfer plays in metabolic modularity and environmental adaptation.

Carbon isotope signatures are routinely used to assess the relative importance of the CBB and rTCA cycles in contemporary and past natural environments, and to infer the key organisms responsible for primary production^75–78^. Although stable isotope signatures may accurately reflect the relative importance of distinct carbon fixation pathways in environmental samples, our study shows that assigning these key ecological functions to particular microbial groups requires a deeper understanding of how the underlying metabolic pathways are distributed in nature.

## Material & Methods

### Sample collection

*“B.” childressi* individuals were collected at cold seeps in the northern Gulf of Mexico at the GC246 and GC234 sites during the R/V Atlantis AT26-13 cruise in April 2014, Nautilus cruise NA044 in July 2014 and Nautilus NA058 cruise in May 2015. The *B. azoricus* individual was collected at the Lucky Strike hydrothermal vent field on the North Mid-Atlantic Ridge (NMAR) during the Biobaz cruise in 2013. A list of samples and fixation details are summarized in Table S7.

### DNA and RNA extraction

DNA was extracted from mussel gill tissue according to Zhou *et al.* (1996)^79^ with the following modifications: An initial overnight incubation step was performed at 37 °C in 360 μl of extraction buffer (100 mM Tris-HCl [pH 8.0], 100 mM sodium EDTA [pH 8.0], 100 mM sodium phosphate [pH 8.0], 1.5 M NaCl, 1% CTAB) and 40 μl of proteinase K (10 mg/ml). For transcriptome sequencing, RNA was extracted with an Allprep(R) DNA/RNA micro kit (Qiagen, Hilden, Germany) according to the manufacturer’s instructions. Concentrations of DNA and RNA were measured with a Qubit^®^ 2.0 Fluorometer (Invitrogen, Eugen, USA).

### Metagenome sequencing and assembly

DNA extracted from gill tissues of one *“B.” childressi* individual was sequenced at the Center for Biotechnology at the University of Bielefeld (Bielefeld, Germany). A total of 471,459,598 paired-end reads (150 bp) and 7,739,150 paired-end reads (250 bp long) were generated on Illumina HiSeq 1500 and MiSeq machines, respectively. DNA extracted from gill tissues of one *“B.” childressi* and one *B. azoricus* individual was sequenced by the Max Planck Genome Center (Cologne, Germany) and generated respectively 57,172,785 and 159,408,731 paired-end reads (150 bp long) on an Illumina HiSeq 2500.

We screened the metagenomic and metatranscriptomic libraries for the presence of campylobacterotal 16S rRNA sequences. The PhyloFlash 2.0 suite (https://github.com/HRGV/phyloFlash) was used to perform RNA small subunit (SSU) screening and reconstructions.

Metagenome assembly was performed as follows: First the raw reads were quality trimmed (Q=2) and Illumina adapters were removed using BBduk (BBmap suite v37.9 from Bushnell B. - sourceforge.net/projects/bbmap/). An initial assembly was performed with Megahit^80^ using default settings. The resulting assembly file was then analyzed with metawatt V2.0 binning tools^81^, and draft genome bins were generated by analyzing contig tetranucleotide frequency, differential coverage and GC content. Contigs belonging to bins with an Campylobacterota taxonomic signature were extracted. The quality-trimmed metagenomic reads were then mapped against the Campylobacterota contigs using Bbmap (BBmap suite v37.9), filtering reads with a minimum identity of 98%. The mapped reads were then used for a new assembly using SPAdes 3.4.2^82^ with default settings. Additional details on the assembly process of “*Ca.* T. azoricus” are described in Supplementary note 3. The bin of the free-living Campylobacterota carrying CBB cycle genes was obtained from the Manus Basin metagenome “NSu-F5” as described in^23^ with three rounds of read-mapping, re-assembly and binning for final bin completion of 92% and 11.7% contamination.

Bin quality was checked with CheckM^45^ and a new iteration of taxonomic binning, mapping and assembly was performed until no contamination from other bacterial strains or host remained in the assembly. Contigs smaller than 900 bp were included in BLAST analysis but excluded from subsequent analyses because they were unlikely to have any relevant genetic information. Genomes were annotated with RAST and cross-checked with IMG ER web servers^83–85^. Genome average nucleotide identity (ANI) and average amino acid identity (AAI) were calculated using the AAI and ANI calculator from the enveomics collection^86^ with the default settings. The specific coverage for genomes and gene was calculated using BBmap.

Raw data was uploaded to the European Nucleotide Archive under the accession numbers: PRJEB19882, PRJEB23284, and PRJEB23286.

### Transcriptome sequencing and processing

Transcriptomes of three *“B.” childressi* individuals were sequenced at the Max Planck Genome Center (Cologne, Germany) details are in Table S7. Transcriptome reads were processed as in Rubin-Blum et al.^87^. Briefly, raw reads were mapped against the “*Ca.* T. childressi” draft genome with BBmap (BBmap suite v.37.09): reads were quality trimmed (Q=2), Illumina adapters removed and a minimum similarity of 98% used to map to the reference genome. The number of transcriptome reads mapping to each gene was estimated with featureCounts v1.5.2^88^. To compare the transcriptome libraries of each individual, a normalization factor was estimated with calcNormFactors based on the trimmed mean of M-values (TMM) implemented in the edgeR version 3.16.5^89^. The TMM normalized read counts were converted to reads per kilobases of exon per million reads mapped (RPKM) with edgeR (http://www.bioconductor.org).

### rTCA cycle gene screening

To confirm presence or absence of the rTCA cycle in the metagenomic and transcriptomic libraries, we created a BLAST database containing published amino acid sequences of Campylobacterota rTCA key genes, citrate lyase, 2-oxoglutarate ferredoxin oxidoreductase and pyruvate ferredoxin kinase. The first metagenomic assembly iterations, as well as the final Campylobacterota bins, were screened using BLASTX against the respective database to detect the presence of potential rTCA cycle genes.

### Phylogenomic reconstruction

Phylogenomic trees were calculated using Phylogenomics-tools (Brandon Seah, https://github.com/kbseah/phylogenomics-tools). The draft genomes “*Ca.* T. childressi” and “*Ca.* T. azoricus” and the free-living Campylobacterotum from Manus basin were compared to the genomes of 41 Campylobacterota representatives. Five Deltaproteobacteria genomes were used as outgroup. Universal marker proteins conserved across all bacteria were screened using Amphora2^90^. Genes present in one copy in every draft genome were selected for the phylogenomic reconstruction (*rpsI, rplT, rpsB, rplM, rpsS, rplK, rplL, frr, rplP, rplA, rplB, pyrG, rpsM, smpB*). Each gene set was aligned using MUSCLE^91^. The concat_align.pl script (phylogenomics-tools) was used for determining the best protein substitution model of each marker alignment (*rpsI ::*LG*, rplT ::*LG*, rpsB::*LG *, rplM::*LG *, rpsS::*LG *, rplK::*RTREV *, rplL::*LG *, frr ::*LG*, rplP ::*LG*, rplA::*LG *, rplB::*LG *, pyrG::*LG *,rpsM::*LG *, smpB::*LG). To calculate the multi-gene phylogeny, the marker genes from each genome were concatenated. The best tree with SH-like aLRT support value was calculated with RAxML^92^ using the tree_calculations.pl script (phylogenomics-tools).

### Phylogenetic analysis

The IMG ER pipeline detected genes with a gammaproteobacterial signature based on homologies to sequences in its database. We extracted and analyzed these sequences with the Geneious software version v 9.1.8^93^ (http://www.geneious.com). Genes predicted by automated annotations were manually verified and curated using the public databases NCBI, Uniprot and Swissprot. Sequences of interest were compared to the NCBI nucleotide and amino acid databases using nucleotide- and amino acid-BLAST. We retrieved closely-related sequences from the BLASTX results on the NCBI non-redundant database. Additionally, other reference sequences were included in the analysis and all sequences were aligned using MUSCLE (v3.6.)^91^. To detect the best substitution model to use for phylogenetic reconstruction, we used the ProtTest3 package^94^ (Model summarized in Supplementary Table 8). Phylogenetic analyses were then performed using Bayesian and Maximum likelihood analyses. Bayesian analysis was performed with MrBayes (v3.2)^95^ under a General Time Reversible model with the best-fitted substitution model. Analyses were performed for two million generations using four parallel Monte Carlo Markov chains. Sample trees were taken every 1000 generations. Maximum likelihood trees were calculated with PHYML^96^ using the best-fitted substitution model. We used 1000 bootstraps as support values for nodes in the trees.

### Codon usage analysis

The codon usage of “*Ca.* T. azoricus” and “*Ca.* T. childressi” genes was determined with CodonW^53^ using default parameters. The Principal Component Analysis was plotted with R (version 3.4.0).

### Bulk isotope analysis

Parts of *“B.” childressi* gill tissues were used for bulk stable isotope analysis. Tissue pieces were oven-dried overnight and ground to a fine powder. The dried tissue was weighed and samples (0.3-0.7 mg dry weight) were packaged in tin capsules for mass spectrometry, and analyzed using a Costech (Valencia, CA USA) elemental analyzer interfaced with a continuous flow Micromass (Manchester, UK) Isoprime isotope ratio mass spectrometer (EA-IRMS) for ^15^N/^14^N and ^13^C/^12^C ratios. Measurements are reported in δ notation [per mil (‰) units] and ovalbumin was used as a routine standard. Precision for δ^13^C and δ^15^N was ± 0.2 ‰ and ± 0.4 ‰.

### Protein extraction and peptide preparation

Parts of the gills (see Supplementary Table 7) of three *“B.” childressi* specimen were used to prepare tryptic digests following the filter-aided sample preparation (FASP) protocol of Wisniewski et al.^97^ with minor modifications^55^. Prior to FASP, cells were disrupted by beat-beating samples in SDT lysis buffer (4% (w/v) SDS, 100 mM Tris-HCl pH 7.6, 0.1 M DTT) using lysing matrix D tubes (MP Biomedicals) before heating to 95 °C for 10 minutes.

To allow binding of peptides to the SCX column for 2D-LC methods, peptides were desalted using Sep-Pak C18 Plus Light Cartridges (Waters) according to the manufacturer’s instructions. A centrifugal vacuum concentrator was used to exchange acetonitrile after peptide elution with 0.2% (v/v) formic acid. The Pierce Micro BCA assay (Thermo Scientific) was used to determine peptide concentrations, following the manufacturer’s instructions.

### 1D and 2D LC-MS/MS

All three samples were analyzed by 1D-LC-MS/MS and 2D-LC-MS/MS as described in Kleiner et al.^39^. Briefly, sample analysis via 1D-LC-MS/MS was run twice. An UltiMate^TM^ 3000 RSLCnano Liquid Chromatograph (Thermo Fisher Scientific) was used to load 1.5-3 μg peptide with loading solvent A (2% acetonitrile, 0.05% trifluoroacetic acid) onto a 5 mm, 300 μm ID C18 Acclaim^®^ PepMap100 pre-column (Thermo Fisher Scientific). Peptides were eluted from the pre-column onto a 50 cm × 75 μm analytical EASY-Spray column packed with PepMap RSLC C18, 2 μm material (Thermo Fisher Scientific) heated to 45 °C. An Easy-Spray source connected the analytical column to a Q Exactive Plus hybrid quadrupole-Orbitrap mass spectrometer (Thermo Fisher Scientific). Separation of peptides on the analytical column was achieved at a flow rate of 225 nl min^−1^ using a 460 min gradient going from 98% buffer A (0.1% formic acid) to 31% buffer B (0.1% formic acid, 80% acetonitrile) in 363 min, then to 50% B in 70 min, to 99% B in 1 min and ending with 99% B. Electrospray ionization (ESI) was used to ionize eluting peptides. Carryover was reduced by two wash runs (injection of 20 μl acetonitrile, 99% eluent B) and one blank run between samples. Data acquisition with the Q Exactive Plus was done as in ^98^.

The 2D-LC-MS/MS experiments were performed as described by Kleiner et al.^39^ with the modification that pH plugs instead of NaCl salt plugs were used for peptide elution from the SCX column. Briefly, 4.5 μg of peptide were loaded with loading solvent B (2% acetonitrile, 0.5% formic acid) onto a 10 cm, 300 μm ID Poros 10 S SCX column (Thermo Fisher Scientific) at a flow rate of 5 μl min^−1^ using the same LC as for 1D-LC-MS/MS. Peptides that did not bind to the SCX column were captured by the C18 pre-column (same as for 1D-LC), which was in-line downstream of the SCX column. The C18 pre-column was then switched in-line with the 50 cm × 75 μm analytical column (same as for 1D) and the breakthrough separated using a gradient of eluent A and B (2% B to 31% B in 82 min, 50% B in 10 min, 99% B in 1 min, holding 99% B for 7 min, back to 2% B in 1 min, holding 2% B for 19 min). Peptides were eluted step-wise from the SCX to the C18 pre-column by injecting 20 μl of pH buffers with increasing pH (pH 2.5-pH 8, CTIBiphase buffers, Column Technology Inc.) from the autosampler. After each pH plug, the C18 pre-column was again switched in-line with the analytical column and peptides separated as above. Between samples, the SCX column was washed twice (injection of 20 μl 4 M NaCl in loading solvent B, 100% eluent B), the RP column once (injection of 20 μl acetonitrile, 99% eluent B) and a blank run was done to reduce carryover. Data was acquired with the Q Exactive Plus as in ^98^.

### Protein identification and quantification

A database containing protein sequences predicted from the metatranscriptomic and - genomic data of the *“B.” childressi* symbiosis generated in this study was used for protein identification as described in the ‘Metagenome assembly’ section above. The cRAP protein sequence database (http://www.thegpm.org/crap/), which contains sequences of common lab contaminants, was appended to the database. The final database contained 38,418 protein sequences. For protein identification, MS/MS spectra were searched against this database using the Sequest HT node in Proteome Discoverer version 2.0.0.802 (Thermo Fisher Scientific) as in^33^.

To quantify proteins, normalized spectral abundance factors (NSAFs)^99^ were calculated per species and multiplied by 100, to give the relative protein abundance in %. For biomass calculations, the method described by Kleiner et al.^39^ was used. Calculations of NSAFs and biomass for each sample were based on the combined data from both 1D-LC-MS/MS runs and the one 2D-LC-MS/MS run.

### Direct Protein-SIF

Stable carbon isotope fingerprints (SIFs) for *“B.” childressi* and its symbionts were determined as described by Kleiner et al.^33^. Human hair with a known δ^13^C value was used as a reference to correct for instrument fractionation. A tryptic digest of the reference material was prepared as described above and with the same 1D-LC-MS/MS method as the samples. Due to the low abundance of the “*Ca.* Thiobarba” symbiont in terms of biomass the six 1D-LC-MS/MS datasets (technical replicate runs of three gill samples) were combined in one peptide identification search to obtain enough peptides for SIF estimation. For peptide identification, MS/MS spectra were searched against the database using the Sequest HT node in Proteome Discoverer version 2.0.0.802 (Thermo Fisher Scientific) and peptide spectral matches were filtered using the Percolator node as described by Petersen et al.^98^. The peptide-spectrum match (PSM) files generated by Proteome Discoverer were exported in tab-delimited text format. The 1D-LC-MS/MS raw files were converted to mzML format using the MSConvertGUI available in the ProteoWizard tool suite^100^. Only the MS^1^ spectra were retained in the mzML files and the spectra were converted to centroided data by Vendor algorithm peak picking. The PSM and mzML files were used as input for the Calis-p software (https://sourceforge.net/projects/calis-p/) to extract peptide isotope distributions and to compute the direct Protein-SIF δ^13^C value for each species^33^. The direct Protein-SIF δ^13^C values were corrected for instrument fragmentation by applying the offset determined by comparing the direct Protein-SIF δ^13^C value of the reference material with its known δ^13^C value.

### Data availability

The metagenomic and metatranscriptomic raw reads are available in the European Nucleotide Archive under Study Accession Number: ERZ772703, PRJEB23286, PRJEB23284 and PRJEB19882.

## Acknowledgements

We would like to thank the crew and captains of the scientific vessels of the following cruises: Atlantis cruise AT26-13, Nautilus cruises NA044 and NA058 and the “Pourquoi Pas?” Biobaz cruise. We also thank the Max Planck Genome Center in Cologne for the genome and transcriptome sequencing. We are grateful to Lizbeth Sayavedra for transcriptome assembly advice and support. We also thank Brandon Seah for his help with the naming of “*Ca*. Thiobarba”. We thank Marc Strous for access to proteomics equipment. The purchase of the proteomics equipment was supported by a grant of the Canadian Foundation for Innovation to Marc Strous.

This work was funded by the Max Planck Society, the DFG Cluster of Excellence ‘The Ocean in the Earth System’ at MARUM (University of Bremen), a European Research Council Advanced Grant (BathyBiome, Grant 340535) and a Gordon and Betty Moore Foundation Marine Microbiology Initiative Investigator Award through Grant GBMF3811 to ND, the European Union (EU) Marie Curie Actions Initial Training Network (ITN) SYMBIOMICS (contract number 264774) and a NSF award #0801741 to Dr. Samantha Joye to fund the AT26-13 expedition to the Gulf of Mexico. JMP was supported by the Vienna Science and Technology Fund (WWTF) through project VRG14-021. MK was supported by the Natural Sciences and Engineering Research Council (NSERC) of Canada through a Banting fellowship and the NC State Chancellor’s Faculty Excellence Program Cluster on Microbiomes and Complex Microbial Communities. TH was supported through a fellowship of the German Academic Exchange Service DAAD.

## Author contributions

A.A., N.L., J.P. and N.D. conceived and developed the study. A.A. and N.L analyzed data. A.A. and H.G.V. performed the metagenomic assemblies. A.A. performed transcriptome and genome analyses as well as phylogenetic reconstructions. M.K and T.H performed proteomic analyses. M.K. developed and performed isotopic fingerprinting method. N.L and A.A provided bulk Isotope analysis. N.L provided key support for isotope work and interpretations. A.M. and D.V.M. provided environmental genomic bin. H.E.T. performed genomic sequencing. S.J and M.S provided key biological samples. A.A. and N.L. wrote the manuscript with support from J.P. and N.D. All authors discussed the results and contributed to the final manuscript.

